# Data on the frequency of non-reproductive adults in a cross-cultural sample of small-scale human societies

**DOI:** 10.1101/032318

**Authors:** Cody T. Ross, Paul L. Hooper, Monique Borgerhoff Mulder

## 1 Background

The goal of this brief communication is to report cross-cultural data with relevance to researchers studying co-operative breeding in humans in the context of other non-human mammals. The data are derived from ethnographic research in 36 small-scale human populations/ cultural groups living on 5 continents.

A number of recent accounts have described *Homo sapiens* as a species that practices cooperative breeding [e.g., Hill and Hurtado (2009); Hrdy (2009); Kramer (2010); Mace and Sear (2005); van Schaik and Burkart (2010)]. These claims raise two important questions: first, do humans in general, or humans under a specific set of conditions, exhibit behaviors conforming to the technical definition of cooperative breeding? And second, to what extent are patterns of behavior and reproduction in humans similar to, or distinct from those found in non-human animals that are classified as cooperative breeders? These questions has been addressed in part through individual empirical cases studies (Hagen and Barrett, 2009; Hill and Hurtado, 2009; Kramer, 2005; Meehan et al, 2013; Strassmann, 2011) and theoretical papers (Burkart et al, 2009; Smaldino et al, 2013). They remain difficult to answer rigorously, however, without cross-cultural data (Hill et al, 2011; Kramer, 2010). While cooperative breeding is defined as adults exhibiting costly behaviors that typically increase the fitness of other adults, a likely consequence of such behavior is likely to be a pool of non-reproductive adults. No studies to our knowledge have brought standardized cross-cultural data on the actual frequency of non-reproductive adults to bear on the topic. We provide such data in this brief communication.

We note that our data report not percentage of nonbreeders, but rather individuals with a measure of zero for various RS proxies (living children, children surviving to age 5, children surviving to age 21, etc.), and as such could reflect a number of different factors (such as: infertility, failure to keep children alive to 5th—or 21st—birthday, or reproduction suppression) that may, or may not, reflect cooperative breeding.

## 2 Methods

For purposes unrelated to questions concerning cooperative breeding in humans, we have compiled a large number of data sets on cross-cultural human reproductive outcomes, that may nevertheless be of interest to researchers studying cooperative breeding. Here, we outline our methods of: 1) data-set construction, and 2) Bayesian meta-analysis of the sex- and group-specific probably of remaining non-reproductive until a given age (we run the model for ages 25 and 45).

### 2.1 Data

Our data sets are wide ranging both spatially and temporally. For data to be included, we require that data collection be based on either total censuses or randomized samples of the population. For example, we do not consider data sets where only the reproductive outcomes of household heads are represented, as this could strongly bias our estimates of the frequency on non-reproductive adults. We detail the site-specific data collection methodology by field-site in Section 4.

### 2.2 Bayesian Meta-Analysis

In each population/cultural group, *j*, we count the total number of males and females with age ≥ 25 in model 1 (and age *j*, 45 in model 2), 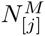 and 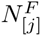, respectively. We then count total number of males and females in this subset that have *RS* = 0, 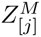 and 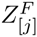Following the multi-level meta-analysis model outlined in Ross et al (2015), we adopt a model where each population/cultural group has its own binomial probability distribution for remaining non-reproductive until age 25 in model 1 (and age 45 in model 2); we describe this probability distribution using the posterior distributions of parameters 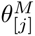 and 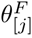, which are unique to males and females respectively. These parameters are estimated using the population-specific count data and multi-level priors:

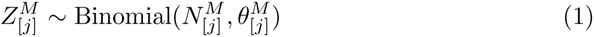

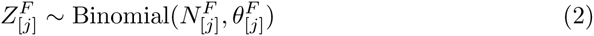

It should be noted that the posterior variances of 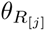 and 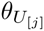, decrease as the sample size in a given population/cultural group increases, which automatically augments the weight that each population/cultural group carries on the estimation of higher order parameters. We utilize the inverse logit function to transform the probabilities, 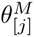 and 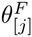, into their respective log-odds expressions, 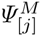 and 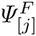, and declare that 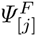 can be described as plus some deviation, 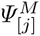, which represents the estimated change in the log-odds of remaining non-reproductive as a function of sex, unique to each subpopulation, *j*.

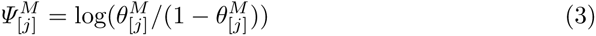

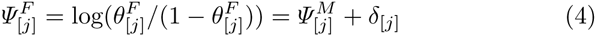

We then model the parameters 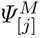 and 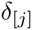 as realizations from higher-level Gaussian distributions, with unknown means and standard deviations, to yield estimates corresponding to the mean log odds across populations of males remaining non-reproductive, and the mean offset in the log odds of males and females remaining non-reproductive:

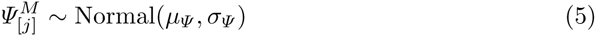

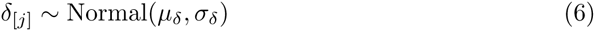

We utilize weakly informative normal priors on *μ_Ψ_* and *μ_δ_*, and weakly informative half-Cauchy priors on *σ_Ψ_* and *σ_δ_* (Gelman et al, 2006):

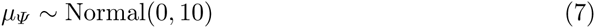

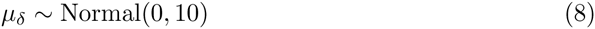

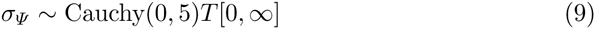

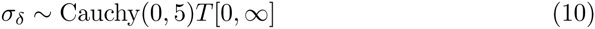

We fit this model using Hamiltonian Monte Carlo, as implemented in the Stan 2.2.0 environment (Stan Development Team, 2013). Two chains were updated adaptively for 1,000 iterations, and then sampled for 3,000 iterations, with no thinning. Stan monitors multiple chain convergence with the 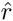 statistic (which equals 1 at convergence), and monitors effective sample size (Gelman and Rubin, 1992). All model parameters had an 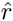 statistic of less than 1.001, and an effective sample size of greater than 2,000. Visual inspection of trace plots showed excellent mixing and apparent convergence of multiple chains to the same equilibrium distribution. Model code and model diagnostics are included in the Supplementary Materials.

## 3 Results

Across study sites, the inferential results suggest that the median percent of non-reproductive adults older than 25 is 0.10 (PCI95: 0.06, 0.15) for males and 0.07 (PCI95: 0.03, 0.13) for females. The corresponding results for adults older than 45 are 0.05 (PCI95: 0.02, 0.09) for males and 0.04 (PCI95: 0.01, 0.09) for females. Figure 1 plots these distributions.

**Fig. 1:**
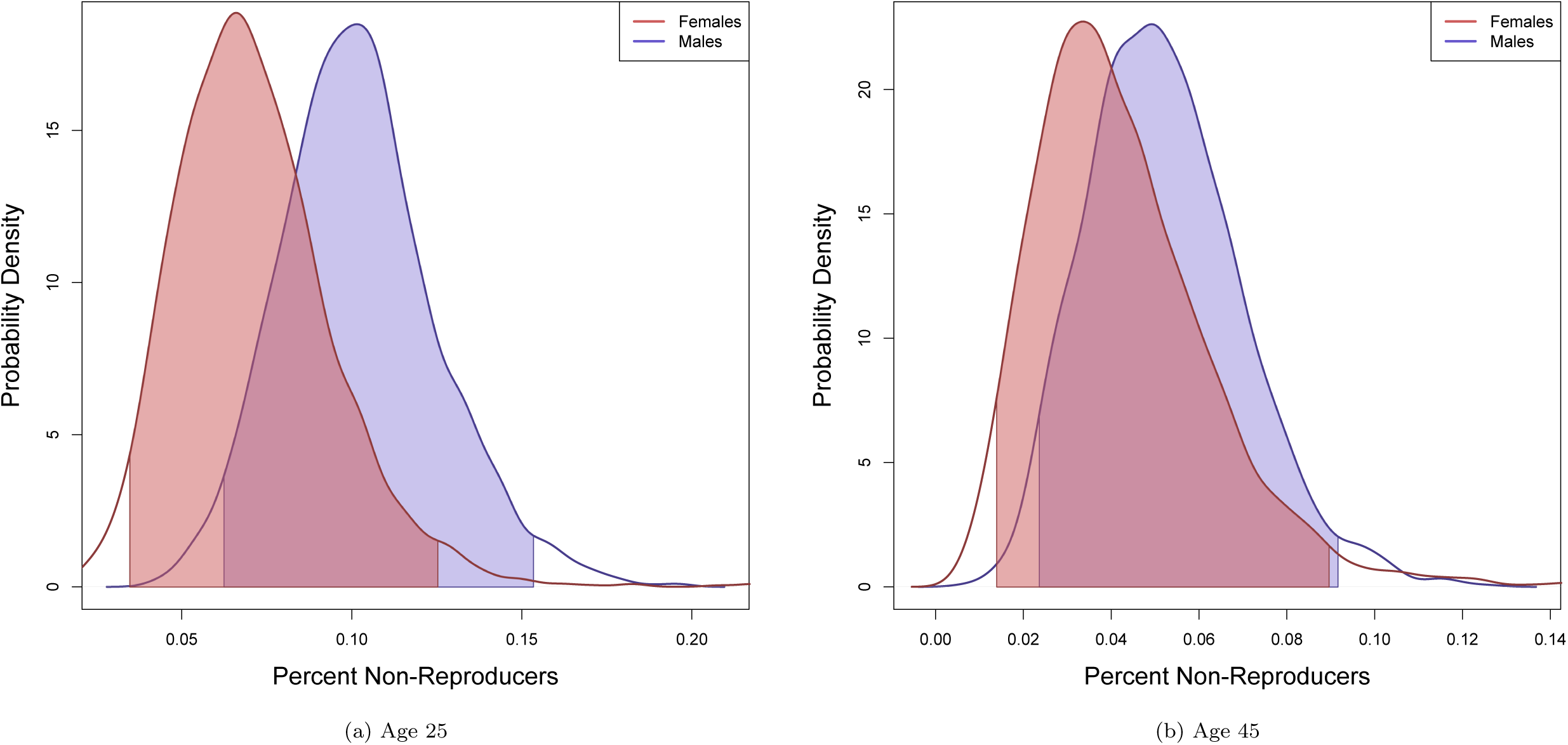
Cross-cultural probability of being non-reproductive at age 25 (Frame 1a) and at age 45 (Frame1b), for males (blue) and females (red).

There is a considerable range in the group- and sex-specific frequency of non-reproductive adults across sites, from near 0.00 (among men and women in a number of Tanzanian samples) to above 0.40 (among Meriam women and English men). Table 1 reports the following summary statistics for females and males at each study site: 1) the total number of adult age ≥ 25, 2) the total number and percent of adults age ≥ 25 who did not produce any surviving offspring, and 3) the posterior estimates from the meta-analysis. Table 2 includes the same estimates but for adults age ≥ 45.

**Table 1:** Frequency of non-reproductive adults age ≥ 25. The symbols 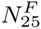 and 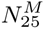 give the sample sizes of males and females over the age of 25 in a given population, respectively. The symbols 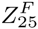 and 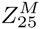 give the corresponding sample sizes of males and females over the age of 25 who have RS=0. The terms *F_pct_* and *M_pct_* give the raw percentages, and the terms *F_post_*, and *M_post_*, give the posterior results from the meta-analysis in terms of a median and a 95 percent posterior confidence region.

| Population | Ethnographer(s) | Location | Subsistence | RS Definition | Method | $N_{25}^F$ | $Z_{25}^F$ | $F_{Pct}^F$ | $F_{Post}^F$ | $N_{25}^M$ | $Z_{25}^M$ | $M_{Pct}$ | $M_{Post}$ | Delta |
| --- | --- | --- | --- | --- | --- | --- | --- | --- | --- | --- | --- | --- | --- | --- |
| Kipsigis | Borgerhoff Mulder | Kenya | Agropastoralism | Survived to 5 | Census | 869 | 16 | 0.02 | 0.02 (0.01, 0.03) | 786 | 110 | 0.14 | 0.14 (0.12, 0.16) | -2 (-2.42, -1.58) |
| Pimbwe | Borgerhoff Mulder | Tanzania | Horticulture | Survived to 5 | Census | 568 | 57 | 0.1 | 0.1 (0.08, 0.12) | 477 | 93 | 0.19 | 0.19 (0.16, 0.22) | -0.75 (-1.05, -0.45) |
| Sukuma[1] | Borgerhoff Mulder | Tanzania | Agropastoralism | Survived to 5 | Sample |  |  |  |  | 63 | 2 | 0.03 | 0.04 (0.01, 0.08) |  |
| Tyva | Hooper | Siberia | Pastoralism | Surviving | Sample | 22 | 0 | 0 | 0.01 (0, 0.04) | 17 | 0 | 0 | 0.02 (0, 0.06) | -0.68 (-2.03, 0.75) |

**Table 2:** Frequency of non-reproductive adults age ≥ 45. The symbols 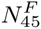 and 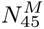give the sample sizes of males and females over the age of 45 in a given population, respectively. The symbols 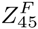 and 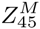, give the corresponding sample sizes of males and females over the age of 45 who have RS=0. The terms *F_pct_* and *M_pct_* give the raw percentages, and the terms *F_post_*, and *M_post_*, give the posterior results from the meta-analysis in terms of a median and a 95 percent posterior confidence region.

| Population | Ethnographer(s) | Location | Subsistence | RS Definition | Method | $N_{25}^F$ | $Z_{25}^F$ | $F_{Pct}^F$ | $F_{Post}^F$ | $N_{25}^M$ | $Z_{25}^M$ | $M_{Pct}$ | $M_{Post}$ | Delta |
| --- | --- | --- | --- | --- | --- | --- | --- | --- | --- | --- | --- | --- | --- | --- |
| Kipsigis | Borgerhoff Mulder | Kenya | Agropastoralism | Survived to 5 | Census | 329 | 5 | 0.02 | 0.02 (0.01, 0.03) | 359 | 55 | 0.15 | 0.15 (0.12, 0.18) | -2.18 (-2.96, -1.57) |
| Pimbwe | Borgerhoff Mulder | Tanzania | Horticulture | Survived to 5 | Census | 211 | 12 | 0.06 | 0.05 (0.03, 0.08) | 194 | 12 | 0.06 | 0.06 (0.03, 0.09) | -0.11 (-0.81, 0.51) |
| Sukuma[1] | Borgerhoff Mulder | Tanzania | Agropastoralism | Survived to 5 | Sample |  |  |  |  | 32 | 0 | 0 | 0.01 (0, 0.04) |  |
| Tyva | Hooper | Siberia | Pastoralism | Surviving | Sample | 21 | 0 | 0 | 0.01 (0, 0.03) | 15 | 0 | 0 | 0.01 (0, 0.05) | -0.61 (-2.54, 1.22) |

Finally, we investigate cross-cultural patterns in sex-specific odds of non-reproduction. For individuals of over 25 years of age, there is a general trend that women have lowered odds of being non-reproducers *μ_δ_* = −0.42 (PCI95: -0.93, 0.06). For individuals over 45 years of age *μ_δ_* = −0.26 (PCI95: -1.09, 0.40), this estimate expands to straddle both sides of zero, indicating that across populations, there can be populations where etiher men or women have increased odds of being non-producers; as before, however, the bulk of this distribution lies on negative values, indicating a general trend that women generally have lowered odds of being non-reproducers. Figure 2 plots the density distributions of these posterior estimates.

**Fig. 2:**
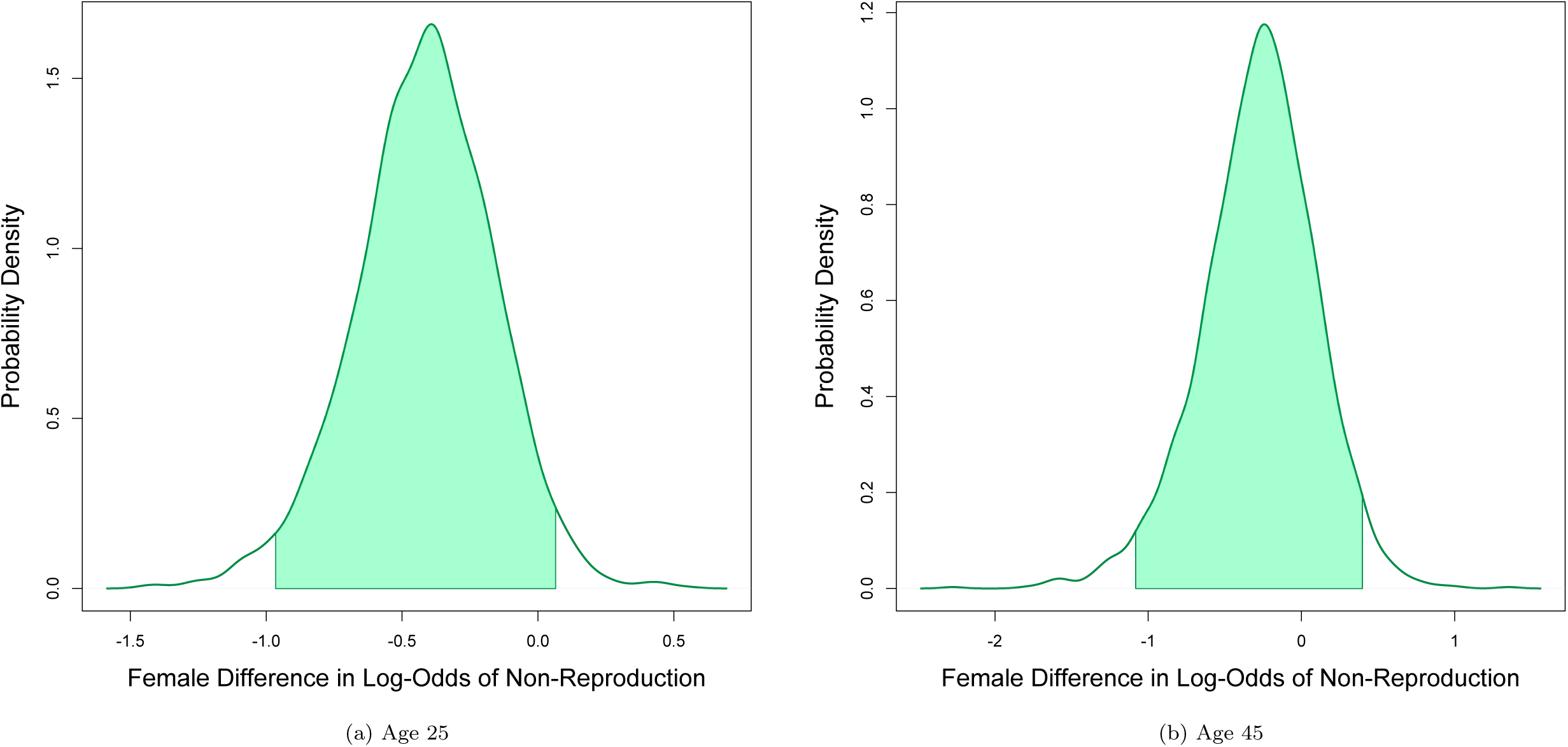
Cross-cultural difference (females relative to males) in log odds of being non-reproductive at age 25 (Frame 2a) and at age 45 (Frame 2b).

## 4 Brief ethnographic information, and data collection methodology by site

### 4.0.1 Kipsigis (Kenya)

The Kipsigis are farming and cattle herding population living in southwestern Kenya (Kericho, Rift Valley Province) (Borgerhoff Mulder, 1987a). They have a strong tradition of polygyny. Residence patterns are strongly virilocal, and the inheritance of land and livestock is strictly patrilineal (Borgerhoff Mulder, 1987b).

The data analyzed here were collected in 1981–1983, with some additional material from 1991. All households, in a cluster of different kokwetinwek (neighbourhoods), were visited and married individuals were interviewed; subsequently unmarried offspring for whom reproductive details were available from their parents were coded into the data set such that all reproductive aged individuals are now sampled. We consider individuals as non-reproductive if they failed to produce any offspring surviving to the age of five years. For younger individuals, age is typically known, but for older individuals a best estimate of age is generated based on *ipinda* (named circumcision age set) membership (Borgerhoff Mulder, 1987a). Full description of the data collection methodology can be found in (Borgerhoff Mulder, 1987a).

### 4.0.2 Pimbwe (Tanzania)

The Pimbwe population studied by Monique Borgerhoff Mulder are a horticultural population living on the north end of the Rukwa Valley in Mpanda District of Western Tanzania; they live off the production of subsistence crops (maize and cassava) and cash crops (maize, sunflower, and occasionally rice) (Borgerhoff Mulder, 2009a,b). They practice serial monogamy (both polyandry and polygyny), although a few men do maintain multiple wives concurrently, but rarely for more than a few years since divorce is common.

The data analyzed here were collected in several panels between 1995 and 2014, and represent a full census of the village of Mirumba (Borgerhoff Mulder, 2009a). To measure reproductive success (*RS*5, offspring born and survived to five years of age), we summed all reported children surviving to age five. See Borgerhoff Mulder (2009a) for further methodological details.

### 4.0.3 Sukuma[2] (Tanzania)

The Sukuma are a group of rapidly expanding agropastoralists in Tanzania (Paciotti and Borgerhoff Mulder, 2004; Paciotti et al, 2005). They are patrilocal and frequently polygynous (Paciotti and Borgerhoff Mulder, 2004). Their subsistence system is based on cattle husbandry and plow farming of rice, maize, peanuts, and potatoes (Paciotti and Borgerhoff Mulder, 2004).

The data analyzed here were opportunistically collected by Monique Borgerhoff Mulder in July-August of 2013. Respondents were typically adult male Sukuma who attended a ward-level seminar on HIV awareness. Follow up interviews were conducted at subsequent collective events, like weddings and village meetings, to get a sample that represents the majority of household heads over 40 years old in the Sukuma community of Kibaoni, Tanzanian and its sub-villages. Each man was interviewed privately. To measure reproductive success *RS*_5_, we summed all reported children surviving to age five.

### 4.0.4 Tyva (Southern Siberia)

Tyva is a mountainous region in Southern Siberia, which was incorporated into the Soviet Union in 1914 and currently lies within the Russian Federation. The traditional economic base of the population of Tyva is semi-nomadic pastoralism based on sheep, goats, cattle, horses, yaks, camels, and reindeer.

The current sample was derived from retrospective demographic interviews collected in 2015 in two pastoral communities of the Bai-Taiga region of western Tyva. Interviews were conducted with all adults born prior to 1965 recording the reproductive histories of their parents, their siblings, and themselves. All individuals in the same generation as the interviewees born prior to 1945 were included in the analysis.

## Acknowledgements

We thank Kim Hill, Joan Silk, and the members of the NIMBioS Working Group on Hierarchy and Leadership in Mammalian Societies for encouragement and helpful input. The data contributors thank their local assistants and the communities with whom they have worked for their cooperation and hospitality. NIMBIos has provide support for this project.

